# Multicenter reverse-phase protein array data integration

**DOI:** 10.1101/2021.08.31.458377

**Authors:** Leanne de Koning, Stephan Bernhardt, Kenneth G. Macleod, Bérengère Ouine, Aurélie Cartier, Vonick Sibut, Neil O. Carragher, Ulrike Korf, Bryan Serrels, Adam Byron

**Affiliations:** Department of Translational Research, Institut Curie, PSL Research University, Paris, France; Division of Molecular Genome Analysis, German Cancer Research Center (DKFZ), Heidelberg, Germany; Cancer Research UK Edinburgh Centre, Institute of Genetics and Cancer, University of Edinburgh, Edinburgh, United Kingdom; U900 INSERM, Institut Curie, PSL Research University, Paris, France

## Abstract

Among the technologies available for protein biomarker discovery and validation, reverse-phase protein array (RPPA) benefits from unequalled sample throughput. Panels of high-quality antibodies enable the quantification by RPPA of protein abundance and posttranslational modifications in biological specimens with high precision and sensitivity. Incorporation of RPPA technology into clinical and drug development pipelines requires robust assays that generate reproducible results across multiple laboratories. We implemented the first international multicenter pilot study to investigate RPPA workflow variability. We characterized the proteomic responses of a series of breast cancer cells to two cancer drugs. This analysis quantified 86,832 sample spots, representing 108 biological samples, arrayed at three independent RPPA platforms. This unique integrated set of data is publicly available as a resource to the proteomic and cancer research communities to catalyse further analysis and investigation. We anticipate that this dataset will form a reference for the comparison of RPPA workflows and reagents, which can be expanded in the future, and will aid the identification of platform-robust treatment-marker antigens in breast cancer cells.

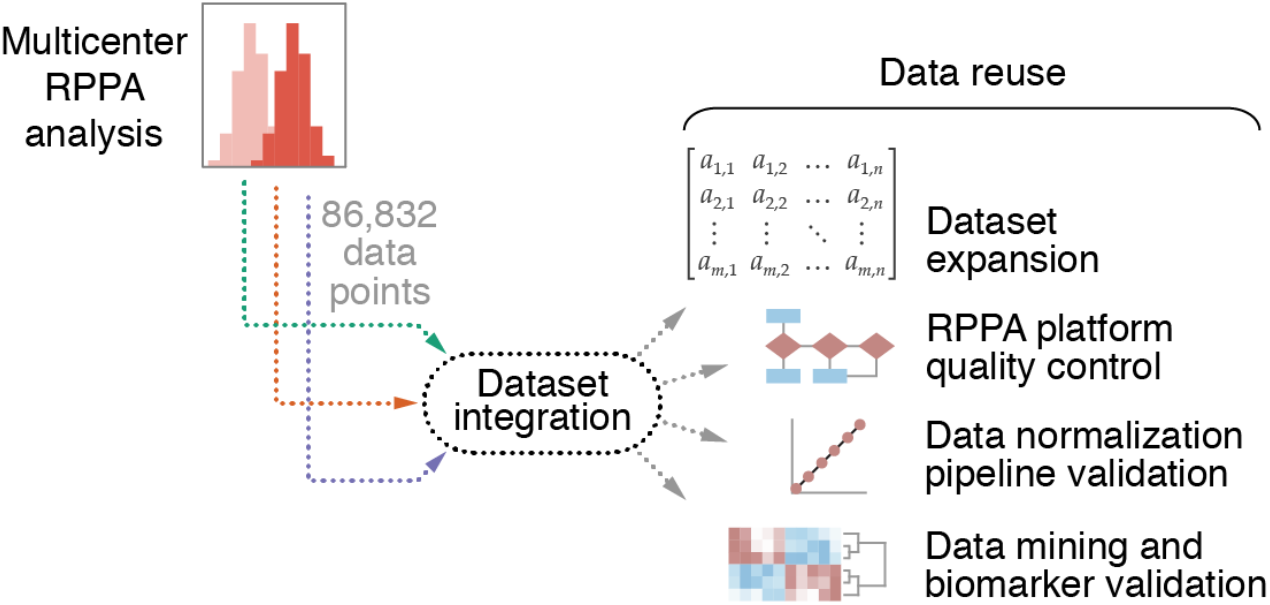

## Background & Summary

In oncology, the therapeutic potential of personalized medicine has been realized for many patients. The targeted therapies imatinib and trastuzumab are examples of drugs that have revolutionized patient survival in certain types of cancer^1,2^. Yet, besides a few success stories, there is little evidence that treatment selection based on genetic alterations is of real benefit to patients^3,4^, possibly because the vast majority of targeted therapies target proteins, and most frequently the active form of the protein. Activation of proteins and their associated signaling pathways often cannot be predicted from genetic alterations. For example, in breast cancer, a *PIK3CA* mutation is not necessarily associated with activation of the phosphoinositide 3-kinase–Akt pathway^5,6^. Moreover, if parallel signaling pathways are activated in the tumor, the patient will show resistance to the administered targeted therapy. An emerging consensus view has therefore identified the need to include proteomic analyses in drug development pipelines and molecular screening programs^7–11^. However, successful implementation of this requires robust, reproducible, and affordable high-throughput proteomics technologies.

Reverse-phase protein array (RPPA) technology is a high-throughput dot-blot technology that uses panels of high-quality antibodies to enable the quantification of protein abundance, and that of posttranslational modifications, from a limited amount of biological sample^12^. The technology offers precise and sensitive multiplexed quantification of the abundance of proteins and posttranslational modifications in hundreds of arrayed samples, providing the opportunity for high-throughput proteomic profiling of cells and tissues^13–15^. RPPA therefore provides a valuable tool for the targeted analysis of disease-relevant signaling mechanisms, including in cancer^16–35^.

To date, RPPA technology has been, and is currently being, applied in several cancer clinical trials^11,36^, mostly retrospectively to detect markers of response and resistance^37–45^, but in some cases also prospectively to select the most appropriate therapy for a given patient^46,47^. As RPPA technology is readying itself for further implementation in drug development and clinical laboratory settings, there was a demand to evaluate the robustness and reproducibility of RPPA datasets derived from different RPPA platforms.

To address this need for crossplatform validation of RPPA technology, we conducted the first international multicenter pilot study to investigate RPPA workflow variability and potential for multiplatform data integration. With the overarching goal of learning from distinct analysis pipelines and improving RPPA technology, the objectives of this open, collaborative project were to acquire, normalize, and integrate data from multiple RPPA platforms to facilitate crossplatform data comparison. We characterized the protein-level responses of six breast cancer cell lines to two clinically relevant cancer drugs at two treatment timepoints, generating a total of 108 biological samples, incorporating 36 experimental conditions, which were normalized and analyzed at three different research centers (Fig. 1a–c). The dataset consists of original RPPA slide scan images and image quantification of 86,832 sample spots derived from 972 arrayed lysates, incorporating independent biological replicate samples, serial lysate dilution series, and technical replicate sample spots, from three independent RPPA platforms. Integrative analysis of a subset of these data enabled the identification of platform-independent protein markers of breast cancer cell response to drug treatment^48^. We report here the raw and processed data from our assessment of multiple RPPA workflows at different research centers. Our data will serve as a starting point for the appraisal of the reproducibility of RPPA technology and its capacity to identify robust protein markers of response to cancer therapies. The data can be reused by multiple researchers over the world to implement and validate their RPPA quality control, data normalization, and data integration methods. Moreover, the dataset can be expanded to support a framework for the integration, interrogation, and interpretation of RPPA data derived from multiple research centers (Fig. 1d).

**Figure 1.**
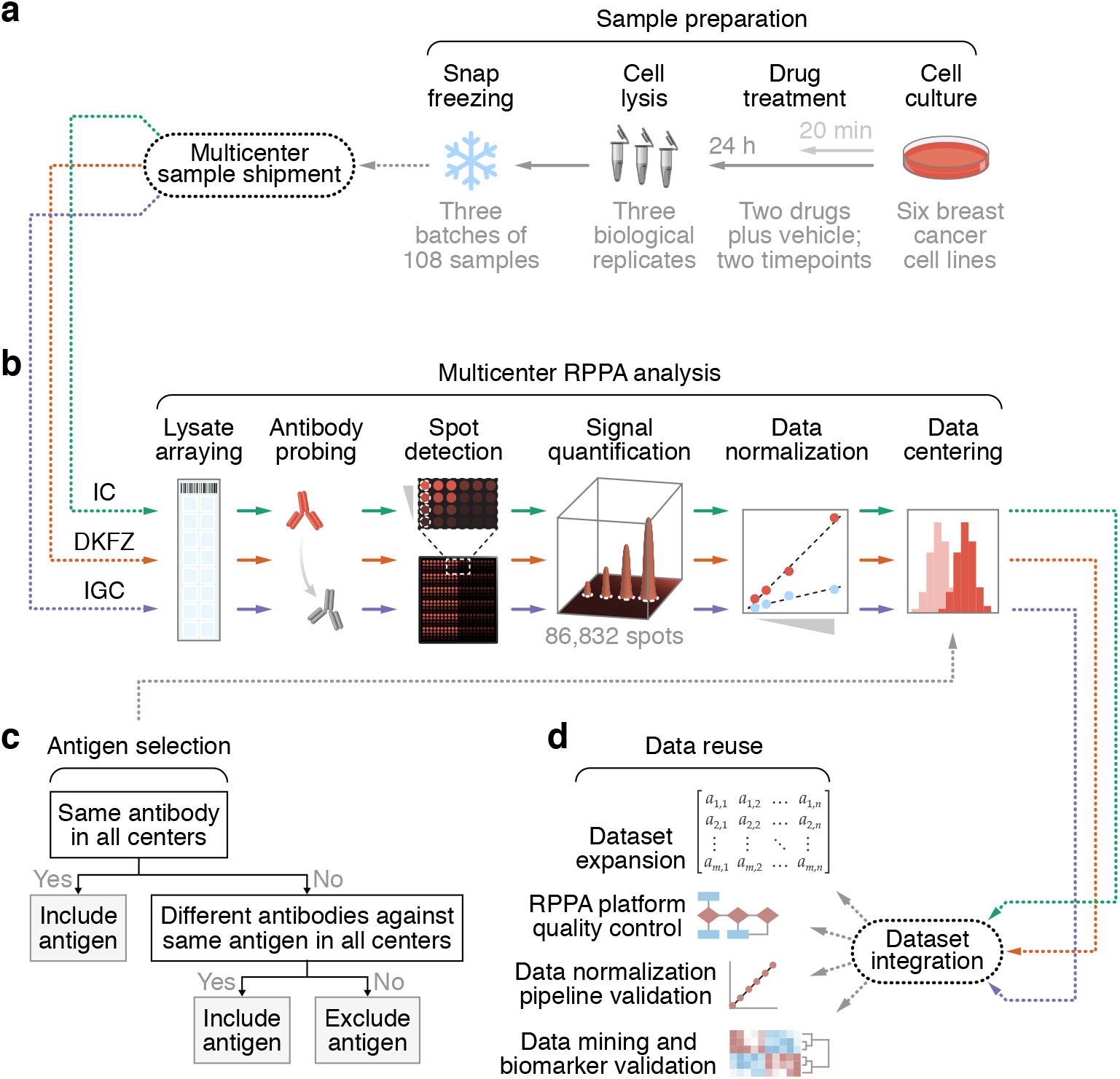
Schematic overview of the sample preparation and data analysis workflows. (**a**) Sample preparation workflow. Six breast cancer cell lines were treated with lapatinib, selumetinib, or DMSO (vehicle) for 20 min or 24 h. Cells were lysed (*n* = 3 biological replicates), and cell lysates were snap frozen and sent to the different research centers. (**b**) Multicenter RPPA analysis workflow. Cell lysates were arrayed and processed for RPPA analysis using the platform-specific setups at the respective research centers [IC, Institut Curie (platform 1); DKFZ, Deutsches Krebsforschungszentrum (platform 2); IGC, Institute of Genetics and Cancer, University of Edinburgh (platform 3)]. (**c**) Antigen selection decision tree. Data derived from validated antibodies targeting the same antigen, including different antibodies from different suppliers, acquired at all three research centers were used for dataset integration. (**d**) Data reuse exemplar workflow. Normalized RPPA data from each research center were centered and integrated to enable various modes of data reuse. a and b were adapted from Fig. 1a in our related work^48^.

## Methods

### Experimental design

For this multicenter RPPA study, six breast cancer cell lines were cultured in the presence or absence of two kinase inhibitors for two time periods. For each experimental condition, three biological replicate cell lysates were generated, frozen, and sent to three different research centers for RPPA analysis (Fig. 1a, b). Cell lysates were processed for RPPA analysis using the platform-specific setups at the respective research centers, including incorporation of serial lysate dilution series and technical replicate sample spots (detailed below). Raw RPPA data were processed and normalized according to the respective in-house data analysis pipelines. Each antibody used was assigned a unique antibody identifier linked to a Research Resource Identifier (RRID) (Online-only Table 1). Normalized data derived from validated antibodies targeting the same antigen acquired at all three research centers were integrated to support data reuse (Fig. 1b–d).

**Table 1.** Breast cancer cell types and drug treatments used to generate samples analyzed in this study. Receptor status of breast cancer cell types is indicated. The breast cancer subtype for MDA-MB-231 cells is also referred to as mesenchymal stem-like. For MDA-MB-453 cells, the *ERBB2* gene, which encodes HER2, is amplified but HER2 is not overexpressed; its breast cancer subtype is sometimes referred to as luminal androgen receptor. E-cad., E-cadherin; EGFR, epidermal growth factor receptor; ER, estrogen receptor; HER2, human epidermal growth factor receptor 2; PR, progesterone receptor.

| Cell type | Breast cancer subtype | Protein expression |  |  |  | *ERBB2* amplification | Drug treatments |
| --- | --- | --- | --- | --- | --- | --- | --- |
|  |  | ER | PR | E-cad. | EGFR-high |  |  |
| HCC1954 | Basal-like HER2-positive | – | – | + | + | + | Lapatinib (20 min, 24 h)<br>Selumetinib (20 min, 24 h)<br>DMSO (20 min, 24 h) |
| MCF7 | Luminal A | + | + | + | – | – | Lapatinib (20 min, 24 h)<br>Selumetinib (20 min, 24 h)<br>DMSO (20 min, 24 h) |
| MDA-MB-231 | Claudin-low | – | – | – | + | – | Lapatinib (20 min, 24 h)<br>Selumetinib (20 min, 24 h)<br>DMSO (20 min, 24 h) |
| MDA-MB-453 | HER2-positive | – | – | – | – | + | Lapatinib (20 min, 24 h)<br>Selumetinib (20 min, 24 h)<br>DMSO (20 min, 24 h) |
| MDA-MB-468 | Basal-like 1 | – | – | + | + | – | Lapatinib (20 min, 24 h)<br>Selumetinib (20 min, 24 h)<br>DMSO (20 min, 24 h) |
| SKBR3 | Luminal-like HER2-positive | – | – | – | + | + | Lapatinib (20 min, 24 h)<br>Selumetinib (20 min, 24 h)<br>DMSO (20 min, 24 h) |

Some of the following methods are expanded versions of descriptions in our related work^48^.

### Cell culture and drug treatment

HCC1954 (RRID CVCL_1259), MCF7 (RRID CVCL_0031), MDA-MB-231 (RRID CVCL_0062), MDA-MB-453 (RRID CVCL_0418), MDA-MB-468 (RRID CVCL_0419), and SKBR3 (RRID CVCL_0033) breast cancer cells (Table 1) were purchased from American Type Culture Collection (LGC Standards, Molsheim, France) and were cultured according to the supplied instructions. Cells were routinely tested for mycoplasma and used within three months of recovery from frozen. Lapatinib [Selleck Chemicals, Houston, TX, USA, catalog number (cat. no.) S2111], a dual epidermal growth factor receptor (EGFR) and human epidermal growth factor receptor 2 (HER2; also known as ErbB2) inhibitor, and selumetinib (Selleck Chemicals, cat. no. S1008), a mitogen-activated protein kinase kinase (MEK) inhibitor, were prepared as 10 mM stock solutions in DMSO (Online-only Table 2). Cells were treated with 1 μM lapatinib, 1 μM selumetinib, or DMSO (vehicle control) in growth medium for 20 min or 24 h.

**Table 2.**
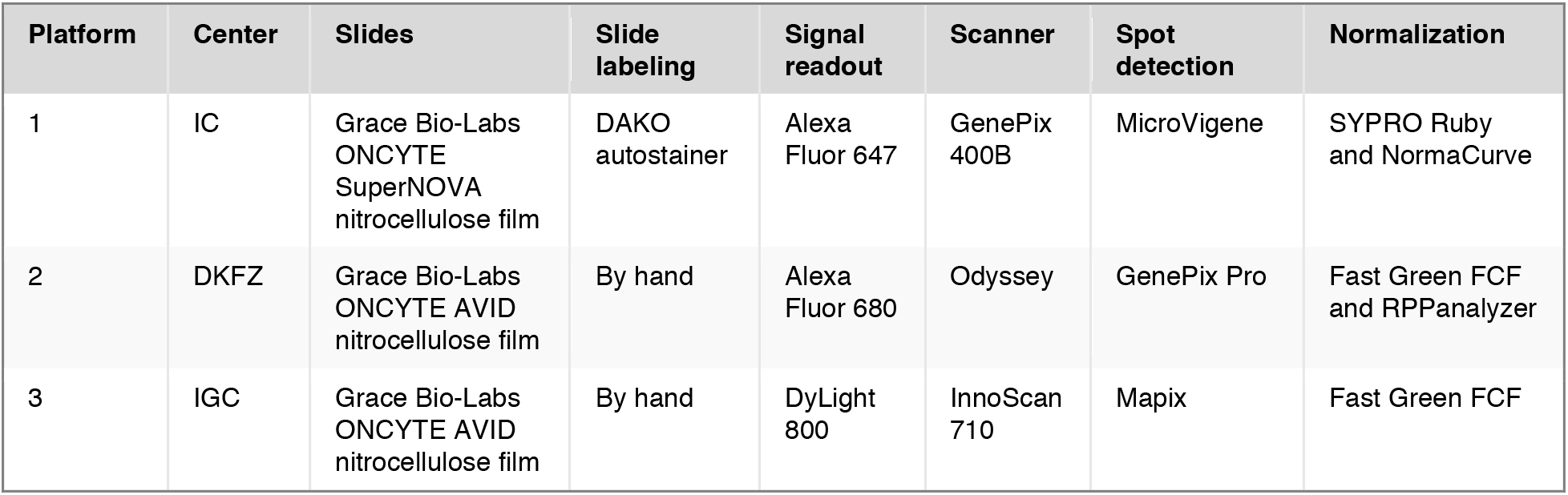
RPPA workflows used at each center. All research centers used the 2470 Arrayer for slide printing. Only platform 1 used signal amplification (Bio-Rad Amplification Reagent). For further details, including manufacturer information, see the Methods section. IC, Institut Curie; DKFZ, Deutsches Krebsforschungszentrum; IGC, Institute of Genetics and Cancer, University of Edinburgh. This table was adapted from Supplementary Table 2 in our related work^48^.

### Cell lysis

Following treatment with drug or vehicle control, cells were washed twice with ice-cold phosphate-buffered saline (PBS). Cells were lysed in hot Laemmli buffer [50 mM Tris-HCl (pH 6.8), 2% (w/v) sodium dodecyl sulfate, 5% (w/v) glycerol, 2 mM DTT, 2.5 mM EDTA, and 2.5 mM EGTA], supplemented with 4 mM sodium orthovanadate, 20 mM sodium fluoride, 1× Perbio Halt phosphatase inhibitor cocktail (Thermo Fisher Scientific, Waltham, MA, USA, cat. no. 78420), and 1× Roche cOmplete protease inhibitor cocktail (Sigma-Aldrich, Gillingham, UK, cat. no. 11697498001), for 10 min at 100°C. Lysates were passed through a 25-gauge needle five times and then clarified by centrifugation at 18,000 × *g* for 10 min at room temperature. Protein concentration was determined for each sample using a Pierce reducing agent-compatible BCA protein assay kit (Thermo Fisher Scientific, cat. no. 23250).

Clarified lysates were aliquoted and snap-frozen in liquid nitrogen prior to shipment to the participating research centers on dry ice. Samples prepared in biological triplicate were analyzed at each research center using the respective in-house RPPA platforms (platform 1, Institut Curie; platform 2, Deutsches Krebsforschungszentrum; platform 3, Institute of Genetics and Cancer, University of Edinburgh) (Table 2).

### RPPA sample handling

#### Platform 1

The protein concentration of all samples was adjusted to 1.0 mg/ml. Samples were then serially diluted in lysis buffer to produce four serial two-fold dilutions of each sample (1 mg/ml, 0.5 mg/ml, 0.25 mg/ml, and 0.125 mg/ml concentrations). Samples were printed onto Grace Bio-Labs ONCYTE SuperNOVA nitrocellulose film slides (Sigma-Aldrich, cat. no. 705170) under conditions of constant 70% relative humidity using an Aushon BioSystems 2470 microarrayer (Quanterix, Billerica, MA, USA). Two technical replicate spots were printed per sample dilution.

#### Platform 2

Vehicle control samples only were serially diluted in lysis buffer to produce six serial two-fold dilutions of each sample (1.8 mg/ml, 0.9 mg/ml, 0.45 mg/ml, 0.225 mg/ml, 0.1125 mg/ml, and 0.05625 mg/ml concentrations). The protein concentration of all samples was then adjusted to 1.5 mg/ml. Samples were printed onto Grace Bio-Labs ONCYTE AVID nitrocellulose film slides (Sigma-Aldrich, cat. no. GBL305108) under conditions of constant 70% relative humidity using an Aushon BioSystems 2470 microarrayer. Three technical replicate spots were printed per sample dilution.

#### Platform 3

The protein concentration of all samples was adjusted to 1.5 mg/ml. Samples were then serially diluted in PBS containing 10% (w/v) glycerol to produce four serial two-fold dilutions of each sample (1.5 mg/ml, 0.75 mg/ml, 0.375 mg/ml, and 0.1875 mg/ml concentrations). Samples were printed onto Grace Bio-Labs ONCYTE AVID nitrocellulose film slides under conditions of constant 70% relative humidity using an Aushon BioSystems 2470 microarrayer. Three technical replicate spots were printed per sample dilution.

### RPPA sample analysis

#### Platform 1

Slides were hydrated in deionized water (four 15-min washes), washed with Tris-buffered saline (TBS) containing 0.1% (w/v) Tween 20 (TBS-T) (three 5-min washes), and incubated with TBS-T containing 5% (w/v) bovine serum albumin (TBS-T-BSA) for 10 min. Slides were washed with TBS-T (three 5-min washes), incubated with Dako biotin (cat. no. X0590) and peroxidase (cat. no. S200389) blocking reagents (both Agilent, Santa Clara, CA, USA), and incubated with primary antibodies (diluted in TBS-T-BSA; Online-only Table 1) for 60 min. Slides were washed with TBS-T (three 5-min washes), incubated with TBS-T-BSA for 10 min, and washed with TBS-T (three 5-min washes). Bound antibodies were detected by incubation with horseradish peroxidase-conjugated antimouse (cat. no. 115-035-062; RRID AB_2338504) or anti-rabbit (cat. no. 111-035-045; RRID AB_2337938) secondary antibodies (both Jackson ImmunoResearch Europe, Ely, UK) for 60 min. To amplify the signal, slides were incubated with Bio-Rad Amplification Reagent (Bio-Rad, Watford, UK, cat. no. 1708230) for 15 min. The arrays were washed with TBS-T (three 5-min washes), probed with Alexa Fluor 647-coupled streptavidin (Thermo Fisher Scientific, cat. no. S32357), incubated with TBS-T-BSA for 60 min, and washed in TBS-T (three 5-min washes). Slides were washed with deionized water (one 5-min wash), spun at 2,000 r.p.m. for 5 min, and stored in the dark.

For normalization against total protein, an additional slide was incubated with destain solution 1 [7% (v/v) acetic acid, 10% (v/v) methanol in deionized water] for 15 min and washed with deionized water (two 5-min washes). The slide was then incubated with SYPRO Ruby staining solution (Thermo Fisher Scientific, cat. no. S11791) for 5 min, washed with deionized water (five 5-min washes), spun at 2,000 r.p.m. for 5 min, and stored in the dark. All steps were performed at room temperature with agitation on a rocking platform.

#### Platform 2

Slides were hydrated in deionized water (four 15-min washes), washed with TBS-T (three 5-min washes), and incubated with 50% (v/v) blocking buffer for fluorescent western blotting (Rockland Immunochemicals, Limerick, PA, USA, cat. no. MB-070) in TBS containing 5 mM sodium fluoride and 1 mM sodium orthovanadate (hereafter, Rockland blocking solution) for 2 h. Slides were washed with TBS-T (four 5-min washes) and incubated with primary antibodies (diluted in Rockland blocking solution; Online-only Table 1) overnight at 4°C with agitation on a rocking platform. Slides were washed with TBS-T (four 5-min washes). Bound antibodies were detected by incubation with Alexa Fluor 680-conjugated F(ab’)_2_ fragments of anti-mouse IgG (cat. no. A-21059; RRID AB_2535725) or anti-rabbit IgG (cat. no. A-21077; RRID AB_2535737) secondary antibodies (both Thermo Fisher Scientific) (diluted 1:8,000 in TBS-T) for 1 h. Slides were washed with TBS-T (four 5-min washes), washed with deionized water (one 5-min wash), air dried for 10 min at 30°C in the dark, and stored at room temperature in the dark.

For normalization against total protein, an additional slide was washed with TBS for 1 min and incubated with 0.005% (w/v) Fast Green FCF (Sigma-Aldrich, cat. no. F7258) in destain solution 2 [10% (v/v) acetic acid, 30% (v/v) ethanol in deionized water] for 45 min. The slide was then incubated with destain solution 2 (two 15-min incubations), washed with deionized water (two 5-min washes), air dried for 10 min at 30°C in the dark, and stored at room temperature in the dark. Unless otherwise stated, all steps were performed at room temperature with agitation on a rocking platform.

#### Platform 3

Slides were hydrated in deionized water (four 15-min washes), incubated with ReBlot Plus strong antigen retrieval agent (Merck, Watford, UK, cat. no. 2504) for 15 min, washed with PBS containing 0.1% (w/v) Tween 20 (PBS-T) (two 5-min washes), and incubated with SuperBlock T20 (TBS) blocking buffer (Thermo Fisher Scientific, cat. no. 37536) for 10 min. Slides were washed with TBS-T (two 5-min washes) and incubated with primary antibodies (diluted in SuperBlock T20 blocking buffer; Online-only Table 1) for 60 min. Slides were washed with TBS-T (two 5-min washes), incubated with SuperBlock T20 blocking buffer for 10 min, and washed with TBS-T (three 5-min washes). Bound antibodies were detected by incubation with DyLight 800-conjugated anti-mouse IgG (cat. no. 5257; RRID AB_10693543) or anti-rabbit IgG (cat. no. 5151; RRID AB_10697505) secondary antibodies (both Cell Signaling Technology) (diluted 1:2,500 in SuperBlock T20 blocking buffer) for 30 min. Slides were washed with TBS-T (two 5-min washes), washed briefly with deionized water, spun at 2,000 r.p.m. for 5 min, and stored in the dark.

For normalization against total protein, an additional slide was washed with deionized water (one 5-min wash), incubated with 1% (w/v) NaOH for 15 min, rinsed with deionized water (10 rinses), washed with deionized water (one 10-min wash), and incubated with destain solution 3 [7% (v/v) acetic acid, 30% (v/v) methanol in deionized water] for 15 min. The slide was then incubated with 0.0025% (w/v) Fast Green FCF in destain solution 3 for 3 min, rinsed with deionized water (10 rinses), incubated with destain solution 3 for 15 min, rinsed with deionized water (10 rinses), spun at 2,000 r.p.m. for 5 min, and stored in the dark. All steps were performed at room temperature with agitation on a rocking platform.

### RPPA data analysis

#### Platform 1

Slides were visualized using a GenePix 4000B microarray scanner (Molecular Devices, San Jose, CA, USA) at an excitation wavelength of 635 nm and a resolution of 10 μm, and images were acquired as tagged image file format (TIFF) files (stored in Images1_IC.zip)^49^. Scanner settings were chosen to obtain around 10–15% of saturated spots, which improves subsequent data normalization. Signals were quantified using MicroVigene microarray image analysis software (VigeneTech, Carlisle, MA, USA). Nonspecific signals were determined by omitting the primary antibody incubation step. Means of technical replicate sample spots were calculated, and data were written to a comma-separated values (CSV) file (SampleQuantification1_IC.csv)^49^. Data were normalized using NormaCurve^50^, which normalizes for fluorescence background per spot and total protein stain and defines a log2-transformed single expression value for each biological sample, based on the entire serial dilution curve and all technical replicates. Next, each RPPA slide was median centered and scaled (divided by median absolute deviation). We then corrected for remaining sample loading effects individually for each array by correcting the dependency of the data for individual arrays on the median value of each sample over all the arrays using linear regression. Normalized, log2-transformed data were written to a CSV file (NormalizedData1_IC.csv)^49^.

#### Platform 2

Slides were visualized using an Odyssey near-infrared microarray scanner (LI-COR Biotechnology, Bad Homburg, Germany) at an excitation wavelength of 685 nm and a resolution of 21 μm, and images were acquired as TIFF files (stored in Images2_DKFZ.zip)^49^. Scanner settings were chosen to enable the highest gain without saturation of the signal. Signals were quantified using GenePix Pro microarray image analysis software (version 7.0) (Molecular Devices). Nonspecific signals were determined by omitting the primary antibody incubation step. The linear fit of the relative fluorescence intensities for the dilution series of each vehicle control-treated sample was determined for each primary antibody, and >99% of signals derived from neat samples (1.5 mg/ml) were within the linear range of detection^48^. Medians of technical replicate sample spots were calculated, background was corrected, and quality control was performed using RPPanalyzer^51^, and data were written to a CSV file (SampleQuantification2_DKFZ.csv)^49^. Data were normalized for protein loading using RPPanalyzer, log2 transformed, and written to a CSV file (NormalizedData2_DKFZ.csv)^49^.

#### Platform 3

Slides were visualized using an InnoScan 710-IR infrared microarray scanner (Innopsys, Carbonne, France) at an excitation wavelength of 785 nm and a resolution of 10 μm, and images were acquired as TIFF files (stored in Images3_IGC.zip)^49^. Following a preview scan to assess signal saturation, scanner settings were chosen to enable the highest laser power and gain with no saturation of the signal. Signals were quantified using Mapix microarray image analysis software (version 6.5.0) (Innopsys). Nonspecific signals were determined by omitting the primary antibody incubation step. The linear fit of the relative fluorescence intensities for the dilution series of each sample was determined for each primary antibody, to assess whether signals were within the linear range of detection with *R*^2^ > 0.9. Mean pixel values of sample spots were corrected by subtracting the median pixel value of the local background surrounding each spot (excluding a 2-pixel region at the spot–background border) using Mapix. Means of technical replicate sample spots were calculated, and data were written to a CSV file (SampleQuantification3_IGC.csv)^49^. Data were normalized for protein loading using the corresponding total protein stain. After data normalization, two data points were negative [cleaved PARP (antibody identifier PARP_cleaved_a) probing of MDA-MB-231 cells treated with vehicle control for 24 h (biological replicate 2) and with selumetinib for 20 min (biological replicate 3)]; to prevent an error upon log2 transformation, these data points were imputed with ~0.98 × column minimum (0.0078125). Imputed, normalized data were log2 transformed and written to a CSV file (NormalizedData3_IGC.csv)^49^.

### Dimensionality reduction

For t-distributed stochastic neighbor embedding (t-SNE), RPPA data were scaled and subjected to principal component analysis, from which the leading 50 principal components were retained for t-SNE. The Barnes–Hut tradeoff parameter, θ, was 0.1, the exaggeration factor was 4, perplexity was 10, and the maximum number of optimization iterations was 5,000. For reproducibility, random seed 1 was used.

For uniform manifold approximation and projection (UMAP), RPPA data were scaled and Euclidean distances between sample points were computed. The local neighborhood used for manifold approximation was 15 neighboring sample points. Fuzzy set operation was 1.0 (pure fuzzy union). Local connectivity was 1, and negative sample rate was 5, with weight γ = 1.0. The effective minimum distance between embedded points was 0.1, and the effective scale of embedded points (spread function) was 1.0. The number of training epochs for optimizing the low-dimensional embedding was 1,000. For reproducibility, random seed 1 was used.

### Multicenter dataset integration

The log_2_ transforms of data normalized using the methods employed by the respective research center were median-subtracted antibody wise and concatenated into a single matrix. To enable comparison across all three research centers, data were filtered for antibodies against the same target (i.e. protein or phosphoprotein) probed at all centers. To consolidate antibody crossreactivity, antibodies recognizing related or multiple protein isoforms (e.g. MEK1 and MEK1/2) were considered to recognize the same target family (e.g. MEK1/2) and were thus included in the integrated dataset if the respective target family was probed at all centers. The final concatenated matrix was written to a CSV file (IntegratedData.csv)^49^.

### Cluster analysis

Agglomerative hierarchical cluster analysis of Spearman rank correlation coefficients was performed using Cluster 3.0 (C Clustering Library, version 1.54)^52^. Spearman rank correlation coefficient-based distance matrices were constructed using pairwise average linkage. Clustering results were visualized using Java TreeView (version 1.1.5r2)^53^.

### Statistical analysis

Three independent biological replicate samples of 36 experimental conditions were analyzed at each research center. No statistical methods were used to predetermine sample size. The median absolute deviation (MAD) was defined as MAD = median[|*x_i_* – median(*x*)|] for a set of biological replicate samples or correlation coefficients *x*_1_, *x*_2_,…, *x_n_*. Robust coefficient of variation based on the median absolute deviation (RCV_*M*_) was defined as RCV_*M*_ = 1.4826 × MAD / median(*x*), where the multiplier 1.4826 is a scaling factor to adjust for asymptotically normal consistency. Pearson correlation coefficient, coefficient of determination, Spearman rank correlation coefficient, and Euclidean distance were calculated for every pairwise combination of biological samples measured by RPPA.

## Data Records

RPPA data records generated at each research center and the integrated RPPA dataset are summarized in Table 3. RPPA slide images generated at each research center are available as TIFF files via figshare^49^. Associated slide descriptor metadata, including antibody position details, and sample descriptor metadata, including spot position details, are available as tab-delimited text files^49^. Sample quantification data files (CSV files) contain background-corrected RPPA intensity data for all biological samples^49^. Normalized data files (CSV files) contain log_2_-transformed data normalized using the methods employed by the respective research center^49^. Centered data files (CSV files) contain antibody-wise median-centered normalized data^49^. The integrated dataset, compiled from the centered data files, is available as a CSV file^49^. A README markdown file (README.md), also available as a plaintext file (README.txt), accompanies the data records^49^.

**Table 3.** Data files generated at each center. A total of 108 biological samples were analyzed at each of the research centers using their in-house RPPA platforms. The dataset comprises RPPA slide scan images and image quantification and normalization of 86,832 sample spots derived from 972 arrayed lysates in total, incorporating independent biological replicate samples, serial lysate dilution series, and technical replicate sample spots, from three independent RPPA platforms. For platform 2, a control dilution series of vehicle control-treated samples only was used (not included in sample spot sum total). Abs, antibodies used; IC, Institut Curie; DKFZ, Deutsches Krebsforschungszentrum; IGC, Institute of Genetics and Cancer, University of Edinburgh.

| Platform | Center | Samples | Serial dilutions | Technical replicates | Abs | Data files | DOI |
| --- | --- | --- | --- | --- | --- | --- | --- |
| 1 | IC | 108 | 4 | 2 | 42 | 61 TIFF files stored in Images1_IC.zip (slide scan images)<sup>49</sup><br>SlideDescriptor1_IC.txt (slide descriptor)<sup>49</sup><br>SampleDescriptor1_IC.txt (sample descriptor)<sup>49</sup><br>SampleQuantification1_IC.csv (sample quantification)<sup>49</sup><br>NormalizedData1_IC.csv (normalized data)<sup>49</sup><br>CenteredData1_IC.csv (centered data)<sup>49</sup> | 10.6084/m9.figshare.14754069<br>10.6084/m9.figshare.14754069<br>10.6084/m9.figshare.14754069<br>10.6084/m9.figshare.14754069<br>10.6084/m9.figshare.14754069<br>10.6084/m9.figshare.14754069 |
| 2 | DKFZ | 108 | 1 | 3 | 44 | 18 TIFF files stored in Images2_DKFZ.zip (slide scan images)<sup>49</sup><br>SlideDescriptor2_DKFZ.txt (slide descriptor)<sup>49</sup><br>SampleDescriptor2_DKFZ.txt (sample descriptor)<sup>49</sup><br>SampleQuantification2_DKFZ.csv (sample quantification)<sup>49</sup><br>NormalizedData2_DKFZ.csv (normalized data)<sup>49</sup><br>CenteredData2_DKFZ.csv (centered data)<sup>49</sup> | 10.6084/m9.figshare.14754069<br>10.6084/m9.figshare.14754069<br>10.6084/m9.figshare.14754069<br>10.6084/m9.figshare.14754069<br>10.6084/m9.figshare.14754069<br>10.6084/m9.figshare.14754069 |
| 3 | IGC | 108 | 4 | 3 | 28 | 17 TIFF files stored in Images3_IGC.zip (slide scan images)<sup>49</sup><br>SlideDescriptor3_IGC.txt (slide descriptor)<sup>49</sup><br>SampleDescriptor3_IGC.txt (sample descriptor)<sup>49</sup><br>SampleQuantification3_IGC.csv (sample quantification)<sup>49</sup><br>NormalizedData3_IGC.csv (normalized data)<sup>49</sup><br>CenteredData3_IGC.csv (centered data)<sup>49</sup> | 10.6084/m9.figshare.14754069<br>10.6084/m9.figshare.14754069<br>10.6084/m9.figshare.14754069<br>10.6084/m9.figshare.14754069<br>10.6084/m9.figshare.14754069<br>10.6084/m9.figshare.14754069 |

**Table 3. Data files generated at each center.**
|  |  |  |  |  |  |  |  |
| --- | --- | --- | --- | --- | --- | --- | --- |
| 1–3 | Integr-<br>ated<br>dataset | 108 | – | – | 53 | IntegratedData.csv (integrated<br>data)<sup>49</sup> | 10.6084/m<br>9.figshare.1<br>4754069 |

## Technical Validation

### Experimental design

Robust, high-quality data are crucial for successful RPPA workflows, especially as RPPA pipelines are continuing to be developed for use in clinical settings^54–58^. Thus, there is a growing need for approaches to record and evaluate the reproducibility of different RPPA platform outputs^59^. To control biological and technical variability in this multicenter study, each cell line was cultured at one center and then lysates were distributed to all centers, so that all platforms analyzed exactly the same batch of protein extract. Sample shipping and transit time have been shown previously to have limited effects on RPPA quantification accuracy^54^. Importantly, the same protein extraction buffer was used for all experiments reported here to avoid differences in efficiency of extraction of nuclear, DNA-bound or membrane-bound proteins. When printing the samples, negative control spots (lysis buffer) were printed in parallel to enable evaluation of background levels. Technical replicate sample spots were printed on different areas of the slides to avoid potential spatial bias.

### Sample variability

The high-throughput nature of RPPA permitted the multiplexed quantification of 108 biological samples at each research center in this study, which yielded quantitative data from a total of 86,832 arrayed sample spots. This large-scale dataset enables the assessment and comparison of variability within and between independent RPPA platforms. Normalized RPPA data for all drug treatments and cell types were similarly distributed for all biological replicate samples (Supplementary Fig. 1). To measure relative variability among biological replicates, robust coefficient of variation based on the median absolute deviation (RCV_*M*_) was calculated using unnormalized RPPA data for all samples across drug treatments (Fig. 2a) and cell types (Fig. 2b). Data from all three RPPA platforms exhibited positively skewed distributions of RCV_*M*_ values, with median RCV_*M*_ ≤ 0.190 for all drug treatments [ranges 0.078–0.185 (platform 1), 0.045–0.063 (platform 2), and 0.098–0.190 (platform 3)] (Fig. 2a). Data for all cell types had median RCV_*M*_ ≤ 0.274 [ranges 0.059–0.274 (platform 1), 0.041–0.107 (platform 2), and 0.076–0.249 (platform 3)], which was no greater than 0.186 if the slightly more variable MDA-MB-231 cell type was excluded (Fig. 2b). Median absolute deviations (MADs) for centered RPPA data from all three RPPA platforms were similarly distributed, with median MADs no greater than 0.153 for all drug treatments (range 0.043–0.153 across platforms) and no greater than 0.211 for all cell types (range 0.037–0.211 across platforms) (Supplementary Fig. 2). These data indicate that biological replicate samples were generally concordant at all research centers.

**Figure 2.**
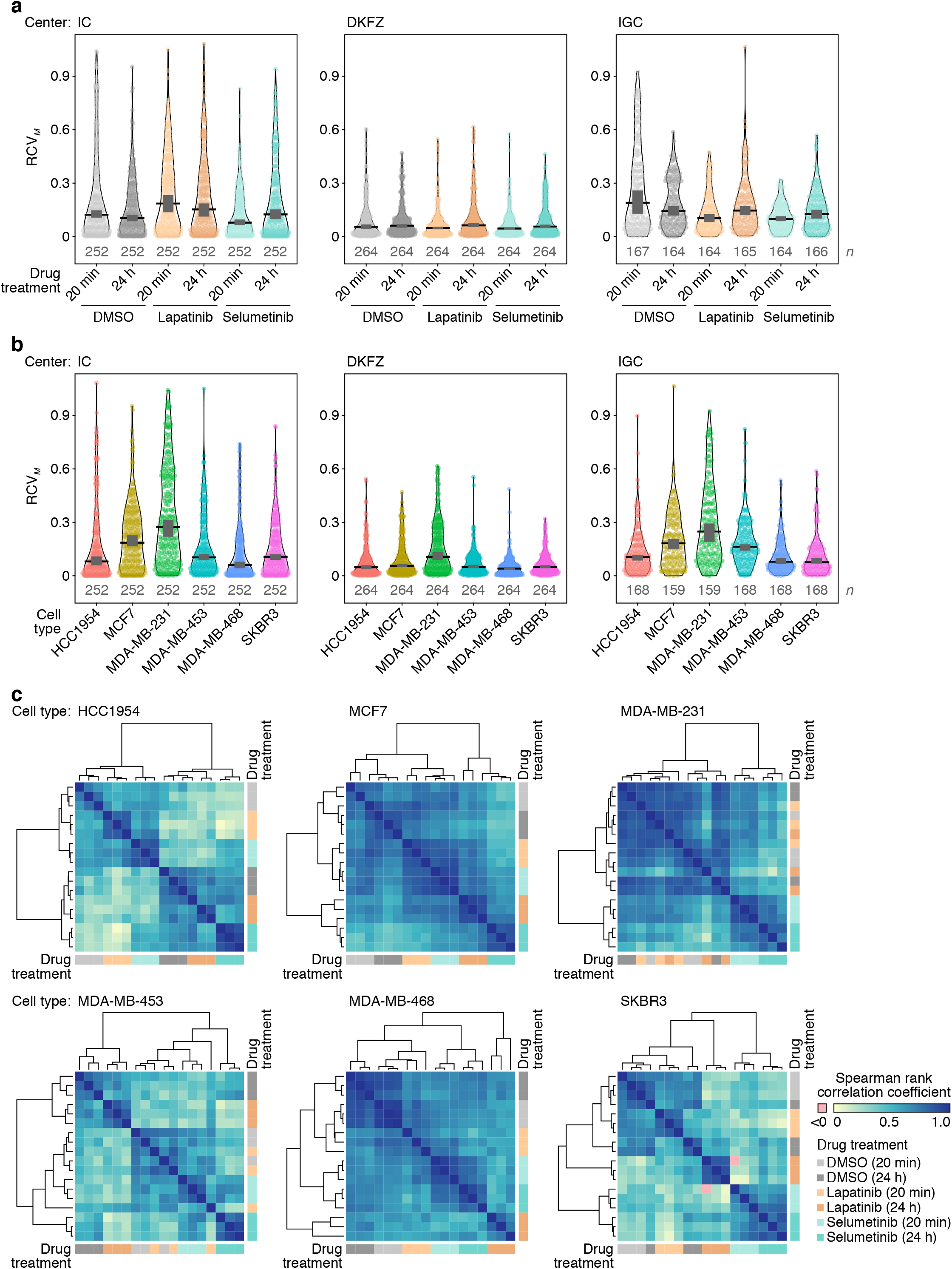
Comparison of RPPA data acquired at each research center. (**a, b**) Distributions of robust coefficient of variation based on the median absolute deviation (RCV_*M*_) for biological replicate sample data derived from all antibodies used at each research center (*n* RCV_*M*_ values, from *n* = 3 biological replicates, are indicated in gray text below each respective probability density). RCV_*M*_ values derived from negative data points were omitted. Unnormalized RPPA data are compared across drug treatments (**a**) and cell types (**b**). Black bar, median; dark gray box, 95% confidence interval; black silhouette outline, probability density. (**c**) Hierarchical cluster analysis of pairwise Spearman rank correlation coefficients between all drug treatments (*n* = 3 biological replicates). Annotation bar color indicates drug treatment and treatment timepoint.

To quantify the extent of agreement between all biological samples tested, multiple similarity metrics were calculated for each of the 11,556 pairwise combinations of RPPA samples (Online-only Table 3). Unsupervised cluster analysis of intersample Spearman rank correlation coefficients for each cell type partitioned sets of biological replicate samples together for most cell types, indicating that they were better correlated than distinct biological samples (Fig. 2c). Indeed, pairwise combinations of like biological replicate samples exhibited strong positive correlations (median ≥ 0.770) with low variability (MAD ≤ 0.066), whereas pairwise combinations of nonreplicate samples were more weakly correlated over a substantially broader range of Spearman rank correlation coefficients (lower limit 0.485 for replicate samples, –0.006 for nonreplicated samples across cell types) (Table 4).

**Table 4.** Correlations between replicate and nonreplicate samples. Pairwise sample Spearman rank correlation coefficients were calculated for like biological replicate samples and for distinct (nonreplicate) samples for each cell type. Minimum (min.) and maximum (max.) values define the range of Spearman rank correlation coefficients for corresponding pairwise sample comparisons.

| Cell type | Replicate sample Spearman rank correlation coefficients |  |  |  |  |  | Nonreplicate sample Spearman rank correlation coefficients |  |  |  |  |
| --- | --- | --- | --- | --- | --- | --- | --- | --- | --- | --- | --- |
|  | Min. | Max. | Median | MAD | *P* |  | Min. | Max. | Median | MAD | *P* |
| HCC1954 | 0.611 | 0.930 | 0.838 | 0.053 | $\leq 3.18 \times 10^{-10}$ | | 0.023 | 0.791 | 0.449 | 0.153 | $\leq 8.35 \times 10^{-1}$ |
| MCF7 | 0.738 | 0.941 | 0.877 | 0.024 | $\leq 3.67 \times 10^{-16}$ | | 0.355 | 0.895 | 0.666 | 0.084 | $\leq 7.41 \times 10^{-4}$ |
| MDA-MB-231 | 0.485 | 0.917 | 0.841 | 0.045 | $\leq 1.88 \times 10^{-6}$ | | 0.126 | 0.920 | 0.680 | 0.131 | $\leq 2.46 \times 10^{-1}$ |
| MDA-MB-453 | 0.500 | 0.925 | 0.770 | 0.046 | $\leq 8.20 \times 10^{-7}$ | | 0.088 | 0.771 | 0.437 | 0.117 | $\leq 4.16 \times 10^{-1}$ |
| MDA-MB-468 | 0.665 | 0.981 | 0.889 | 0.066 | $\leq 2.04 \times 10^{-12}$ | | 0.397 | 0.895 | 0.650 | 0.073 | $\leq 1.43 \times 10^{-4}$ |
| SKBR3 | 0.618 | 0.918 | 0.814 | 0.052 | $\leq 1.74 \times 10^{-10}$ | | -0.006 | 0.819 | 0.414 | 0.122 | $\leq 9.60 \times 10^{-1}$ |

### Cell type-specific RPPA profiles

To explore the protein-level relationships between the different breast cancer cell types examined in this study, we analyzed all centered RPPA data from each research center using t-SNE (Fig. 3a). This dimensionality reduction approach enabled the projection of the high-dimensional RPPA data to two-dimensional space, facilitating the assessment of the structure of each dataset. Visualization using t-SNE projection identified clusters of samples that were partitioned by cell type for each research center (Fig. 3a), supporting the notion that the RPPA data from each platform captured cell type-specific molecular profiles. We confirmed the clustering of cell types in the RPPA data from each platform using an alternative dimensionality reduction approach, UMAP (Supplementary Fig. 3a). In addition to analyses of centered RPPA data, which were used for multicenter dataset integration, we performed t-SNE and UMAP analyses of uncentered normalized RPPA data, which resulted in very similar projections to the centered data (Supplementary Fig. 3b, c), verifying that antibody-wise centering did not distort the data structure.

**Figure 3.**
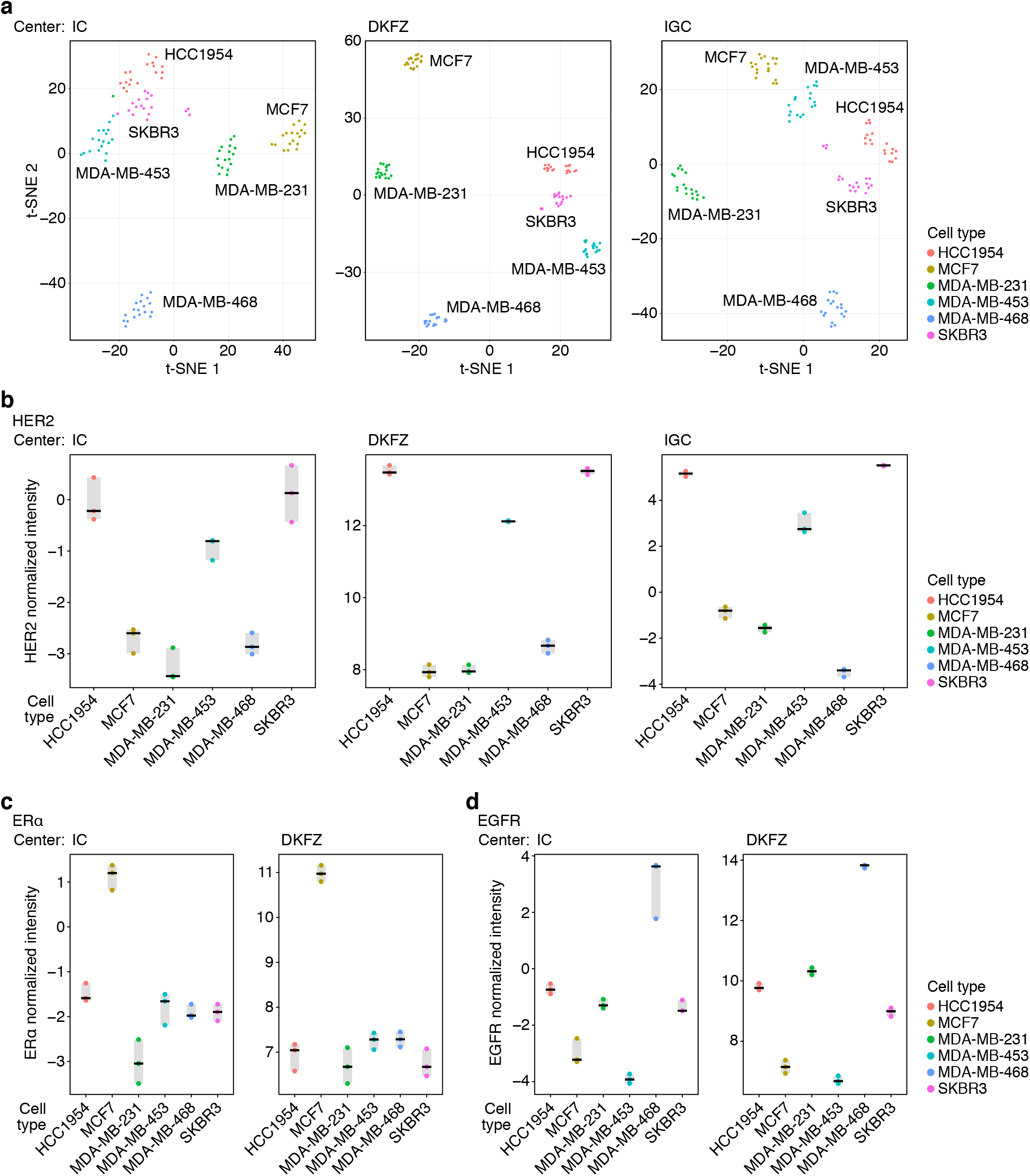
Cell type-specific differences identified by RPPA. (**a**) Dimensionality reduction of centered RPPA data derived from all antibodies used at each research center using t-SNE (*n* = 4,536, 4,752, and 3,024 arrayed lysates for IC, DKFZ, and IGC research centers, respectively, from *n* = 108 biological samples). Cell type classes were colored as indicated. (b–d) Quantification of receptor expression in vehicle control-treated cells (DMSO, 24 h) analyzed by RPPA (*n* = 3 biological replicates). HER2 antibody identifiers for IC, Her2_b; DKFZ, Her2_d; IGC, Her2_a (**b**). ERα antibody identifier for IC and DKFZ, ER-alpha_b (**c**). EGFR antibody identifiers for IC, EGFR_a; DKFZ, EGFR_b (**d**). ERα and EGFR not analyzed at IGC. Black bar, median; light gray box, range.

Samples from some cell types, such as HCC1954 and SKBR3, tended to colocate in t-SNE and UMAP space (Fig. 3a, Supplementary Fig. 3a–c), suggesting that their RPPA profiles were more similar compared to the other cell types. Indeed, the six breast cancer cell lines were chosen to reflect different molecular subtypes of breast cancer (Table 1). Overexpression of the cell-surface receptor HER2 was correctly detected in HCC1954, MDA-MB-453, and SKBR3 cell lines (Fig. 3b); MCF7 was correctly identified as the only estrogen receptor (ER)-positive cell line (Fig. 3c); MDA-MB-468 was correctly identified as the highly EGFR-overexpressing cell line (Fig. 3d). These data imply that cell type-specific protein levels quantified by RPPA can distinguish distinct molecular profiles in breast cancer cell types. Moreover, similar coclustering of these cell types was detected in data from all research centers (Fig. 3a, Supplementary Fig. 3a–c), confirming the capacity for all three RPPA platforms to identify related cell type-specific data structures.

### Cell type-specific drug sensitivities

The panel of breast cancer cell lines examined in this study were treated with two clinically relevant drugs, selumetinib (a MEK inhibitor) and lapatinib (a HER2/EGFR inhibitor) (Online-only Table 2). The sensitivity of these cell lines towards these drugs and the expected changes in major cell signaling pathways are known (http://www.cancerrxgene.org; refs 60–62), which thus served as an internal quality control. We assessed the protein-level responses of the different cell lines following drug treatment. All platforms readily detected the effect of selumetinib on the MEK–extracellular signal-regulated kinase (Erk) pathway, and in particular the blockade between phospho-MEK and phospho-Erk, resulting in the accumulation of phospho-MEK and decrease of phospho-Erk (ref. 63) (Fig. 4a). The two cell lines known to be sensitive to lapatinib (HCC1954 and SKBR3) showed, as expected, downregulation of phospho-HER2 and phospho-EGFR, as well as downregulation of downstream phospho-Akt and phospho-S6 ribosomal protein (Fig. 4b). Cluster analysis of intersample correlations also revealed subclustering of drug-treated samples that was specific to different cell types (Fig. 2c). For example, for MDA-MB-231 cells, Spearman rank correlation coefficients of samples treated with selumetinib (20 min and 24 h) partitioned from the other drug and control treatments, whereas for SKBR3 cells, samples treated with lapatinib (24 h) and selumetinib (20 min and 24 h) partitioned separately from the other treatment conditions (Fig. 2c).

**Figure 4.**
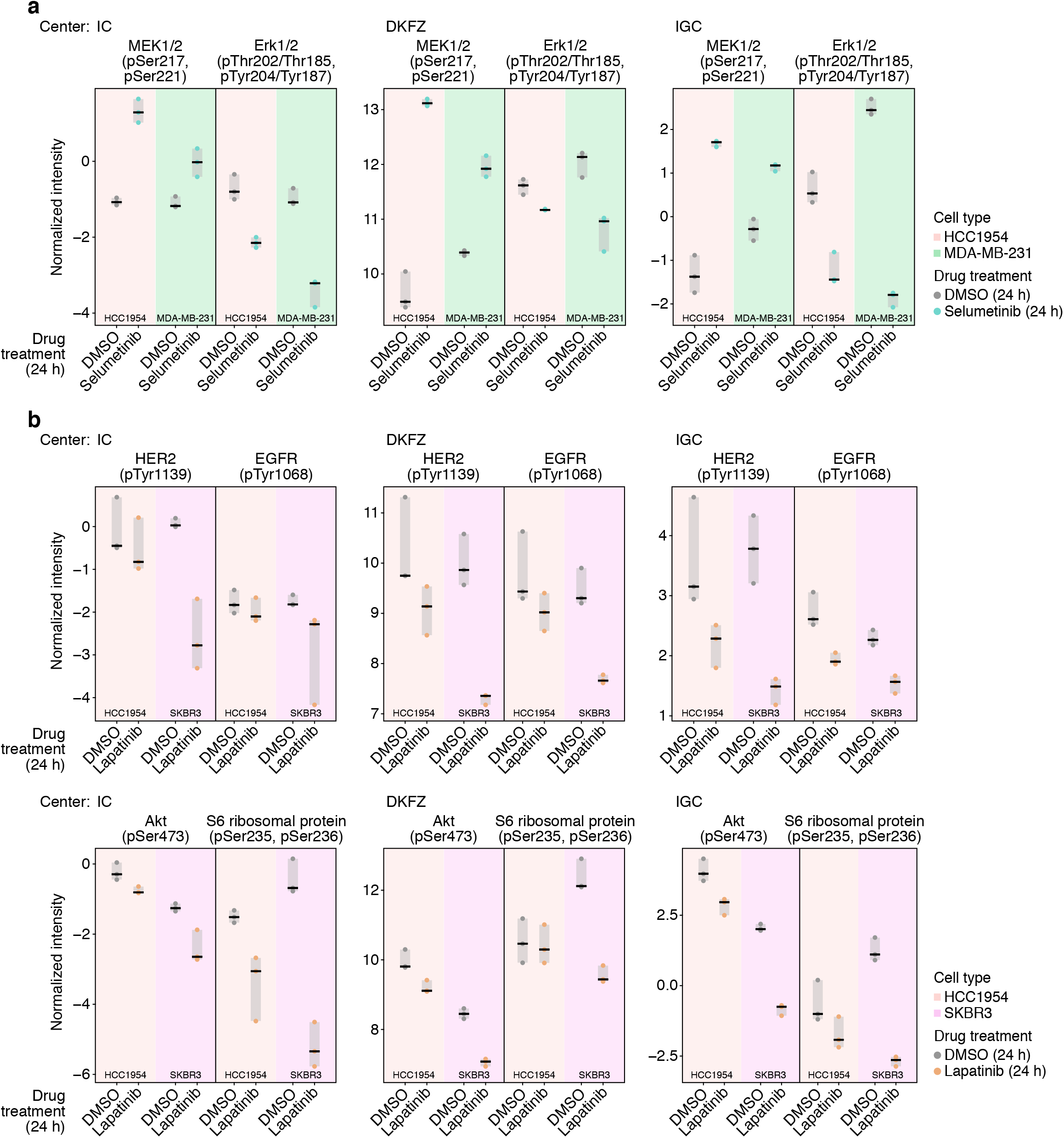
Cell type-specific drug sensitivities identified by RPPA. (**a**) Quantification of phospho-MEK and phospho-Erk in selumetinib- and vehicle control-treated HCC1954 and MDA-MB-231 cells (24 h) analyzed by RPPA (*n* = 3 biological replicates). MEK1/2 (pSer2l7, pSer22l) antibody identifier for IC, DKFZ, and IGC, MEK1/2_pSer217,pSer221. Erk1/2 (pThr2O2/Thr185, pTyr204/Tyr187) antibody identifier for IC, DKFZ, and IGC, Erk1/2_pThr202/Thr185,pTyr204/Tyr187_b. (**b**) Quantification of phospho-HER2, phospho-EGFR, phospho-Akt, and phospho-S6 ribosomal protein in lapatinib- and vehicle control-treated HCC1954 and SKBR3 cells (24 h) analyzed by RPPA (*n* = 3 biological replicates). HER2 (pTyr1139) antibody identifier for IC, DKFZ, and IGC, Her2_pTyr1139. EGFR (pTyr1068) antibody identifiers for IC and IGC, EGFR_pTyr1068_a; DKFZ, EGFR_pTyr1068_b. Akt (pSer473) antibody identifier for IC, DKFZ, and IGC, Akt_pSer473_a. S6 ribosomal protein (pSer235, pSer236) antibody identifiers for IC and IGC, S6 ribosomal protein_pSer235,pSer236_a; DKFZ, S6 ribosomal protein_pSer235,pSer236_b. Black bar, median; light gray box, range.

Together, these data suggest that RPPA data from each platform have the capacity to reveal inherent differences in drug sensitivity and pharmacodynamics between cell types, including those based on their response to targeted therapies.

In summary, the dataset represents high-quality RPPA data derived from multiple distinct RPPA platforms, characterizing a range of human breast cancer cell lines and their response to two clinically relevant cancer drugs. Our collaborative multicenter proteomic approach will enable further dataset integration for the analysis of breast cancer therapeutic response and data reuse for the assessment of interplatform reproducibility.

## Usage Notes

The data reported here serve as a valuable resource for the examination of protein-level responses to pharmacological inhibition in a panel of breast cancer cells, including the search for protein markers of response to clinically relevant therapeutics (Fig. 1d). Such investigations may be facilitated by the interrogation of the highdimensional RPPA data using statistical and machine learning approaches, the functional analysis of signaling pathway changes, or the modeling of protein complexes with protein interaction networks^64^. Indeed, RPPA technology is well placed to elucidate the mechanism-of-action of novel hit compounds or candidate drugs discovered by modern phenotypic screening approaches^65^. Robust and reproducible RPPA platforms are required for profiling large numbers of small-molecule hits from cell-based phenotypic screening assays across doseresponse and time series. Building a repository of publicly available RPPA data associated with pharmacological perturbation, such as that exemplified here, will provide valuable datasets for reuse in emerging machine learning approaches which aim to predict mechanism-of-action or differentiate novel compounds based on RPPA signature similarity with well characterized compounds. This, for example, could extend the Connectivity Map concept – the cataloging and connecting of transcriptional responses to genetic, chemical, and disease perturbations^66^ – by providing additional, orthogonal data at the dynamic posttranslational pathway level.

Moreover, our multicenter analysis provides an important reference for the assessment of RPPA platform robustness and technology reproducibility. Individual RPPA platforms may use the dataset to quality control their workflows, by reproducing the experiments outlined here and comparing their results with our data (Fig. 1d). For interplatform comparison, only antigens analyzed by RPPA at all research centers were compiled in the integrated dataset in this study (Fig. 1c), so future data reuse may include the reintegration of data for antibodies targeting excluded antigens, which are available in the raw data files^49^, to enable analysis of data acquired on different combinations of RPPA platforms. In addition, as no standard tools exist to normalize RPPA data, the raw data reported here may be used to test and validate new data normalization pipelines (Fig. 1d). The data also complement ongoing efforts to assess the effects of sample handling on RPPA quantification in the setting of multisite clinical trials^54^. Finally, the integrated dataset can act as a framework for future interplatform comparisons, and this study opens the door for crossplatform validation of RPPA data to identify robust markers of disease and response to therapy. Expansion of the current dataset to include additional samples, antibodies, or RPPA platforms will improve the applicability of such projects and expand the scope of analysis to, for example, other drug treatments or disease models (Fig. 1d).

The data reported here thus provide a valuable springboard for the assessment of reproducibility or benchmarking of platform-specific RPPA datasets, which will support the refinement of technology best practice and may aid development toward standardization of data processing procedures and improvements in RPPA platform interoperability.

## Code Availability

The code to implement RPPA data analysis using NormaCurve^50^ is freely available at http://microarrays.curie.fr/publications/U900-RPPA_PLT/Normacurve. The code to implement RPPA data analysis using RPPanalyzer^51^ is freely available at https://cran.r-project.org/package=RPPanalyzer. The code to visualize data using PlotsOfData^67^ is freely available at https://github.com/JoachimGoedhart/PlotsOfData. The code to visualize data using UMAP and t-SNE is freely available at https://github.com/atamaianalytics/DimensionReduction.

## Acknowledgments

We thank Stéphane Liva and Patrick Poullet for development and maintenance of data management tools at the Institut Curie research center and Henry Beetham, Philippe Hupé, and Andy Sims for helpful discussions. RPPA platform 1 (Institut Curie research center) is supported by Cancéropôle Ile-de-France. K.G.M. is supported by the Cancer Research UK Edinburgh Centre award (C157/A25140). N.O.C. is supported by Cancer Research UK (C42454/A28596) and the Brain Tumour Charity (GN-000676). A.B. was supported by Cancer Research UK.

## Author Contributions

L.d.K., N.O.C., U.K., B.S., and A.B. conceptualized the study and designed the experiments. S.B., K.G.M., B.O., and A.C. generated the data. L.d.K., S.B., K.G.M., V.S., B.S., and A.B. analyzed the data. A.B. designed and implemented multicenter data integration, curated the data, and prepared the figures. L.d.K. and A.B. and wrote the manuscript. L.d.K., S.B., K.G.M., N.O.C., and A.B. edited the manuscript. All authors critically reviewed and approved the final manuscript.

