## Supplementary Figures for "Multicenter reverse-phase protein array data integration"

**Supplementary Information**

Supplementary Figure 1
Supplementary Figure 2
Supplementary Figure 3
Online-only Table 1
Online-only Table 2
Online-only Table 3

[1]Department of Translational Research, Institut Curie, PSL Research University, Paris, France. [2]Division of Molecular Genome Analysis, German Cancer Research Center (DKFZ), Heidelberg, Germany. [3]Cancer Research UK Edinburgh Centre, Institute of Genetics and Cancer, University of Edinburgh, Edinburgh, United Kingdom. [4]U900 INSERM, Institut Curie, PSL Research University, Paris, France. Present addresses: S.B., Pfizer Pharma GmbH, Berlin, Germany. A.C., Sederma, Le Perray-en-Yvelines, France. B.S., NanoString Technologies, Inc., Seattle, Washington, United States of America. *Correspondence should be addressed to A.B..

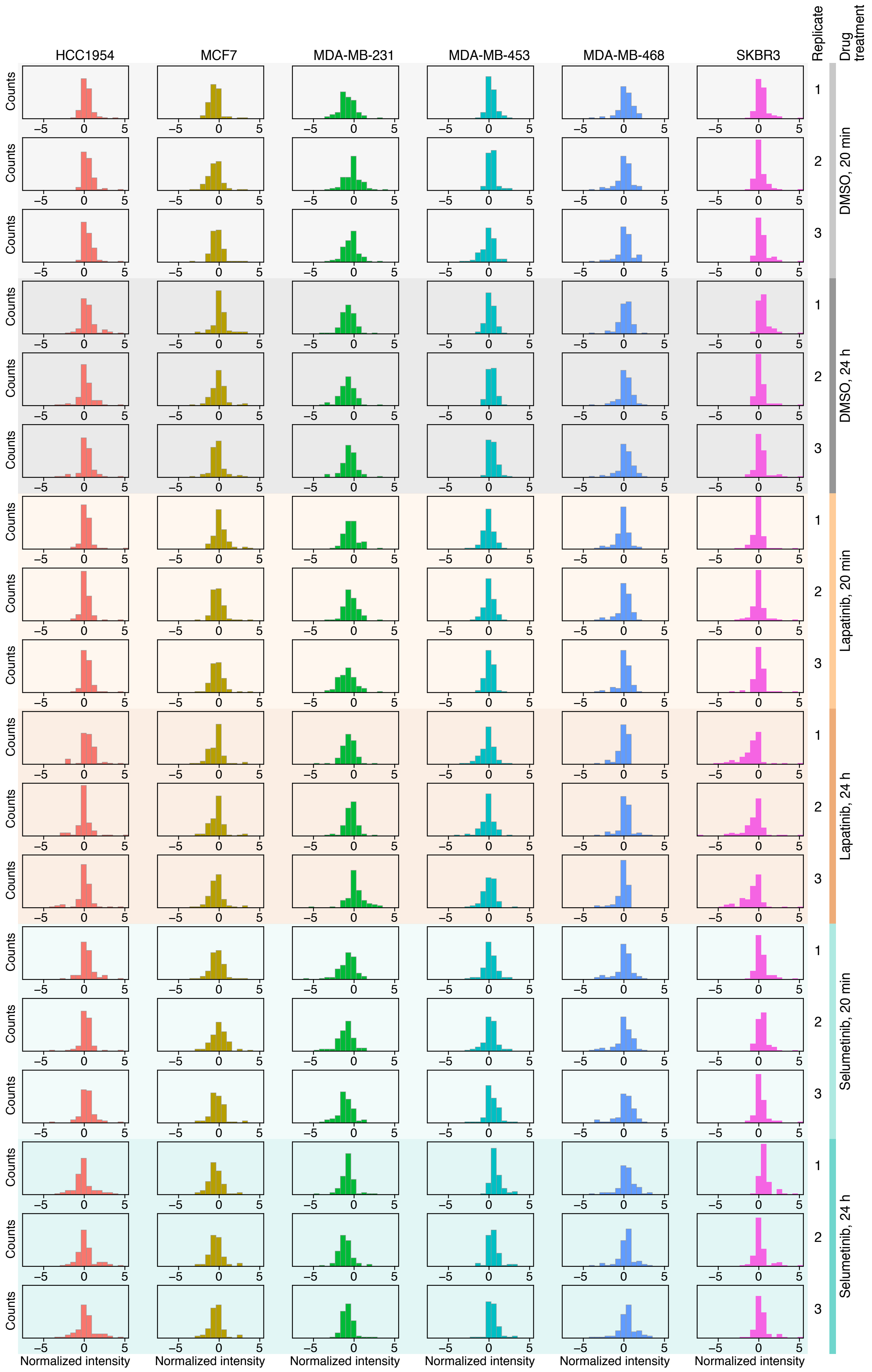


**Supplementary Figure 1.** Distributions of normalized intensities for all samples. Histograms show normalized intensities for all cell types, biological replicates, and drug treatments analyzed by RPPA at all research centers.

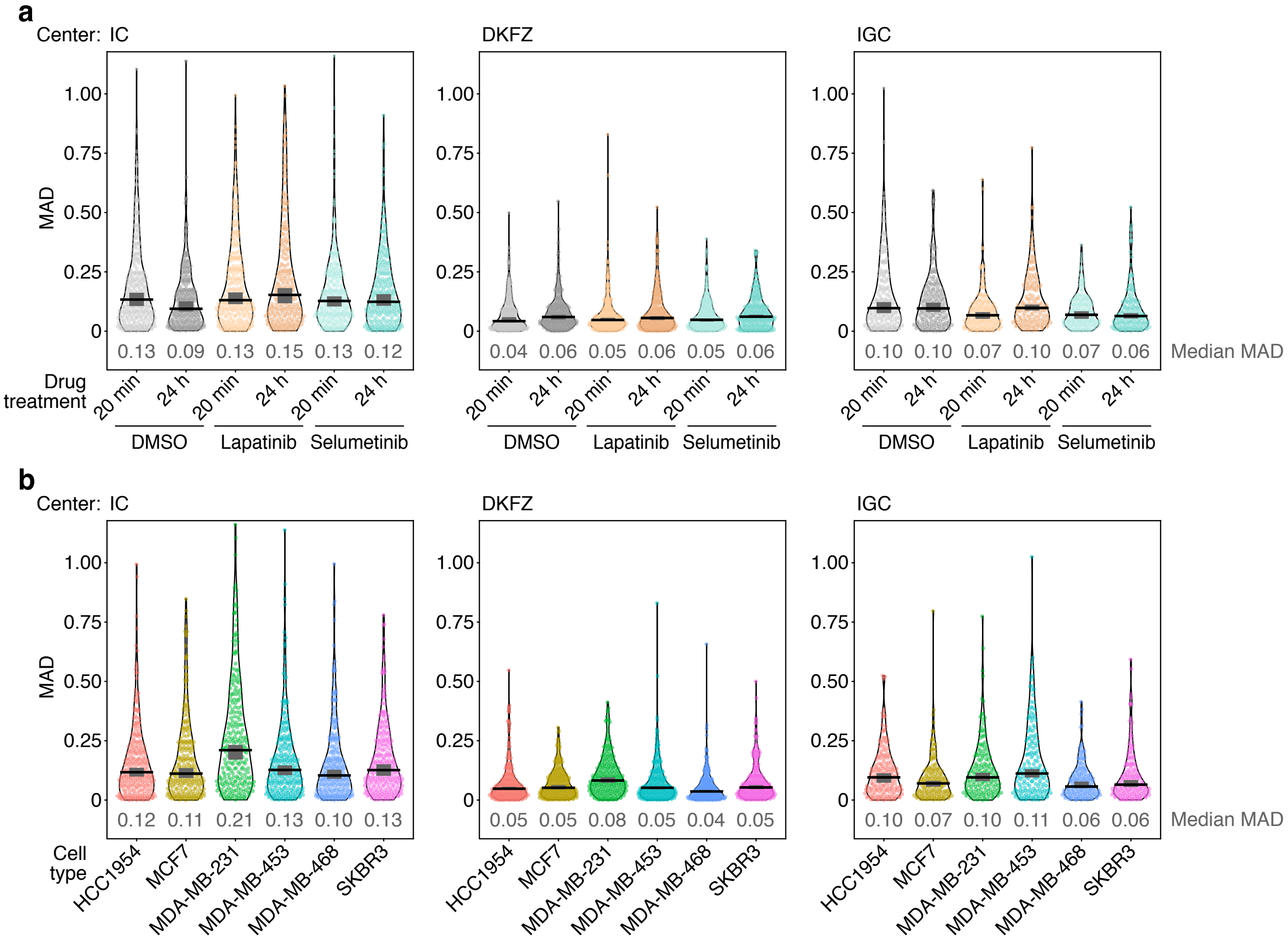


**Supplementary Figure 2.** Evaluation of variability of RPPA data acquired at each research center. (**a**, **b**) Distributions of median absolute deviations (MADs) for biological replicate sample data derived from all antibodies used at each research center ($n$ = 252, 264, and 168 sample MADs for IC, DKFZ, and IGC, respectively, from $n$ = 3 biological replicates). Centered RPPA data are compared across drug treatments (**a**) and cell types (**b**). Black bar, median; dark gray box, 95% confidence interval; black silhouette outline, probability density. Median MAD values are indicated in gray text below each respective probability density.

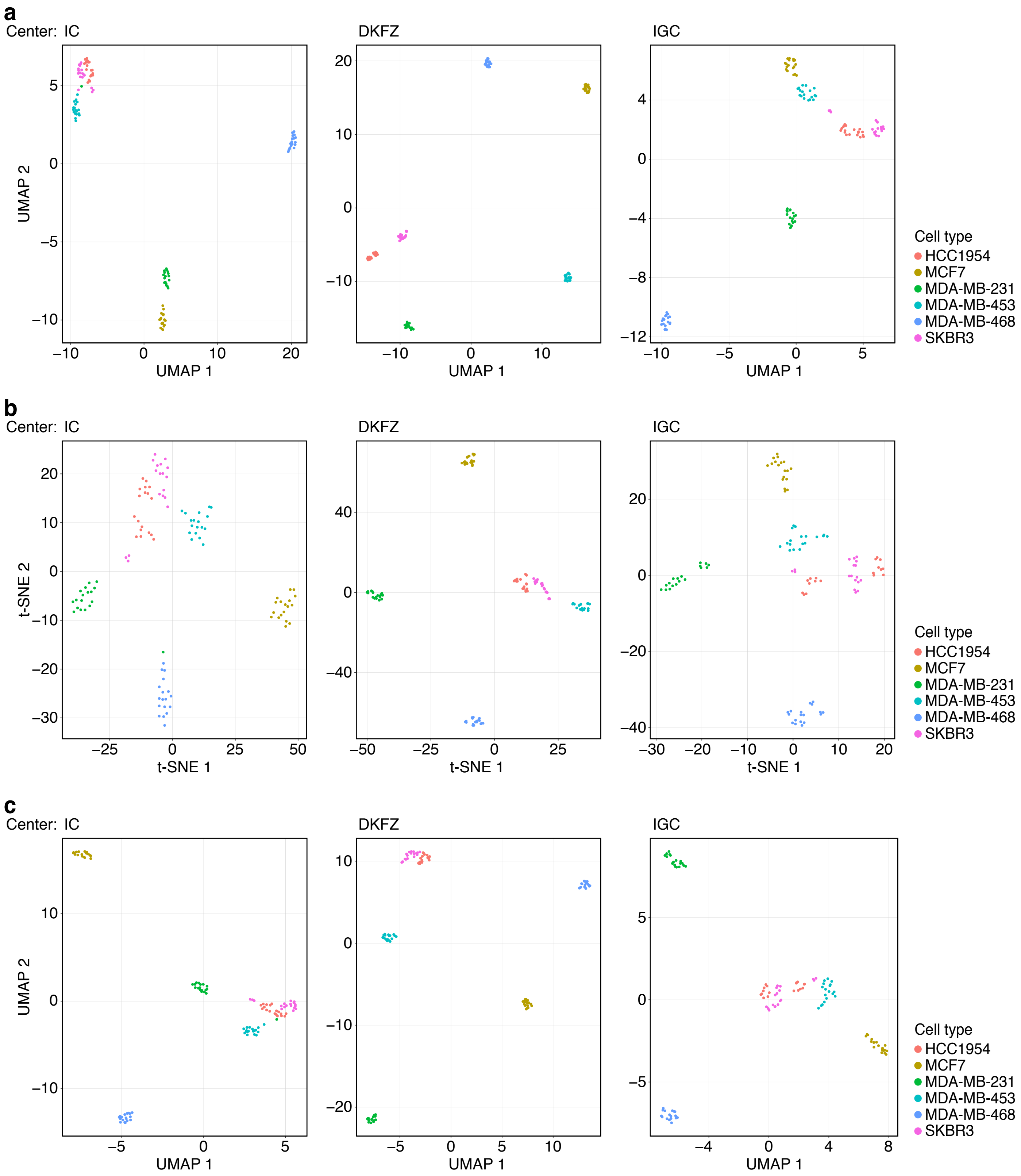


**Supplementary Figure 3.** Visualization of RPPA data acquired at each research center. (**a**) Dimensionality reduction of centered RPPA data derived from all antibodies used at each research center using UMAP. (**b**, **c**) Dimensionality reduction of uncentered RPPA data derived from all antibodies used at each research center using t-SNE (**b**) and UMAP (**c**). For all analyses, $n$ = 4,536, 4,752, and 3,024 sample–antibody combinations for IC, DKFZ, and IGC research centers, respectively, from $n$ = 108 biological samples. Cell type classes were colored as indicated.
